# Single-Cell Analytics for Dose Response (SCADR) discriminates PTEN missense variants by lipid and protein phosphatase dysfunction

**DOI:** 10.64898/2026.08.28.747931

**Authors:** Corbin Glufka, Mahir Taher, Jerry Shijie Tong, Wun Chey Sin, Yue Huang, Patrick Coleman, Fabian Meili, Warren M. Meyers, Paul Pavlidis, Kurt Haas

## Abstract

The proliferation of sequencing efforts has revealed a vast and expanding catalog of single nucleotide gene variants, many associated to, but with unclear roles in disease. Fully charactering variant impacts and linking specific protein dysfunctions to disease are challenging due to the multi-functional nature of many proteins and varying degree of variant effects on these functions. Lagging are sensitive approaches to empirically assess the impact of missense variant-induced single amino acid changes on a wide range of protein functions. To address these issues, we have developed an open-source computational analysis tool called *SCADR* (**S**ingle-**C**ell **A**nalytics for **D**ose **R**esponse) for simultaneously measuring and comparing impacts of exogenously-expressed variants on multiple signaling pathways using multiplex phospho-antibody spectral flow cytometry in human cell lines. SCADR retains and correlates single-cell measures of signal protein activity states along with expression levels of exogenously-expressed variants, providing rich characterization of multiple protein functions, signaling protein interactions, and enhanced discrimination of variant impacts on different signaling pathways, highlighting each variant’s unique dysfunction profile. Here, we apply SCADR for analyses of the impact of 6 variants of the tumor-suppressor protein PTEN (P38H, C124S, G129E, Y138L, D268E, 4A) expressed in HEK293 cells on the phosphorylation states of the canonical and noncanonical downstream signaling proteins Akt, S6, CREB, ERK, and p38 detected with fluorophore-conjugated phospho-antibodies, along with an antibody detecting an N-terminal HA tag on PTEN variants allowing measures of dose-response effects of each variant’s expression on signaling cascades. Results identify variant-specific impacts on downstream signaling cascades.

## INTRODUCTION

Large-scale sequencing efforts have revealed remarkable genetic variation across human populations ^1–5^, and single nucleotide polymorphisms within coding regions that produce missense protein variants have emerged as major contributors to disease ^6–9^, including autism spectrum disorder (ASD; MIM 209850) ^10–12^ and somatic cancers ^13; 14^. However, distinguishing benign from pathogenic variants, and mechanistically linking pathogenic variants to disease, remains difficult given the enormous and continually expanding catalogue of variants identified across human populations. The rate at which new variants are discovered vastly outpaces the field’s capacity to empirically test their effects on protein function and pathophysiology ^15; 16^ and the rarity of most individual variants limits the power of statistical association studies to establish disease links. Computational tools that predict variant pathogenicity help triage this burden ^15^, but in silico estimates remain imperfect, vary in quality from gene to gene, and ultimately require empirical validation. Moreover, most prediction algorithms collapse complex pathogenicity assessments into a single-dimension score, obscuring the multi-faceted nature of protein function and its varied contributions to distinct disease states. Consistent with this concern, wet-lab functional variomic studies have shown that missense variants can disrupt proteins through multiple, distinct molecular mechanisms, effects that may touch all, or only a subset, of a protein’s functions, often in a cell type- or cell state-dependent manner ^7; 17–22^. Needed are approaches capable of rapidly assessing large numbers of variants while remaining sensitive to a broad range of protein functions, a particular necessity for multi-functional proteins, or those whose relationships to disease are still poorly defined.

Here we present an approach, together with custom software, for discriminating variants based on their impact on multiple protein functions, assessed via multiplex single-cell measurements of downstream signaling cascades. Variants are expressed in human cell lines by transfection of expression plasmids, and their effects are probed using fluorophore-conjugated phospho-antibodies that report the activation states of multiple signaling proteins simultaneously. Spectral flow cytometry ^23; 24^ enables the concurrent detection and discrimination of numerous fluorophores within the same cell. Unlike conventional flow cytometry analyses, which typically collapse single-cell measurements into a population-level median fluorescence intensity (MFI) ^25^; ^26^, we introduce a scalable, open-source, high-dimensional analytics platform, **S**ingle-Cell **A**nalytics for **D**ose-**R**esponse (SCADR; github.com/JerryTong-GH/SCADR), that preserves single-cell resolution. SCADR supports single-cell-level correlation analyses, or the binning of cells by exogenous variant-expression level, to extract pathway coupling, expression-dependent signaling effects, and complex, variant-specific functional deficits. By preserving single-cell measurements, or grouping cells according to expression of selected markers, SCADR enables simultaneous tracking of multiple signaling pathways in a manner sensitive to state-dependent pathway interactions, including the dose-dependent effects of exogenously expressed variants.

We demonstrate SCADR’s utility by distinguishing the impact of 6 PTEN (phosphatase and tensin homolog deleted on chromosome ten) variants on canonical and noncanonical signaling pathways, relative to the most commonly expressed NCBI Reference Sequence (RefSeq). PTEN is a dual lipid and protein phosphatase ^27; 28^ whose missense variants have been implicated in somatic cancers ^29^, PTEN hamartoma tumor syndrome (PHTS; MIM 158350) ^30^, ASD ^31^, and other neurodevelopmental disorders. Efforts to link PTEN dysfunction to disease have focused largely on its canonical role regulating the PI3K/Akt/mTOR cell-growth pathway through its lipid phosphatase activity ^20; 32^. However, growing evidence points to additional, noncanonical roles for PTEN, including functions tied to its protein phosphatase activity ^33–36^. Altogether, little is known about how individual PTEN variants affect canonical versus noncanonical signaling, whether a given variant disrupts one or all of PTEN’s functions, or how these distinct dysfunctions relate to the differing pathophysiologies of PTEN-associated diseases.

By combining multiplex signaling-protein profiling with SCADR analytics, we show that subtle impacts of missense variants can be distinguished and their complex relationships to pathway dysfunction characterized. Beyond confirming previously reported effects of well-validated variants on the canonical Akt/S6 pathway, SCADR enabled deeper analysis of their state-dependent effects on signaling-protein interactions. We identified novel impacts of PTEN variants on the noncanonical targets p38, CREB, and ERK: pCREB levels were reduced across the D268E, 4A, Y138L, C124S, and P38H variants relative to RefSeq, and pp38 levels were decreased specifically with Y138L, consistent with a role for PTEN’s protein phosphatase activity in negatively regulating p38. D268E additionally showed reduced pERK levels, indicating a gain-of-function (GoF) effect. Spearman correlation analysis further revealed increased co-regulation among pAkt, pS6, and pp38 in RefSeq and GoF variants, supporting the existence of a PTEN/Akt/p38 signaling axis.

## MATERIALS AND METHODS

### Variant cloning

The PTEN Reference Sequence (RefSeq; NCBI GenBank accession AAB66902.1) and the following 6 PTEN variants were generated by site-directed mutagenesis using Agilent Pfu polymerase, as previously described ^20^, and subcloned using Gateway LR Clonase (Invitrogen) [38] into a eukaryotic expression vector encoding an N-terminal triple hemagglutinin (3×HA) epitope tag: P38H (NM_000314.8:c.113C>A, p.(Pro38His)), C124S (NM_000314.8:c.370T>A, p.(Cys124Ser)). G129E (NM_000314.8:c.386G>A, p.(Gly129Glu)), Y138L (NM_000314.8:c.412_414delTATinsCTG, p.(Tyr138Leu)), D268E (NM_000314.8:c.804C>G, p.(Asp268Glu)), and the synthetic 4A (Ser380Ala;Thr382Ala;Thr383Ala;Ser385Ala).

### Cell culture and transfection

HEK293 cells from the American Type Culture Collection (CRL-1573)^37^ were cultured in Dulbecco’s Modified Eagle’s Medium (Millipore Sigma) supplemented with 10% fetal bovine serum and 100U/mL Penicillin-Streptomycin. Cells were seeded at 2 x10^5^ per well in 24-well plates 16-20 hr before transfection with 1µL of X-tremeGENE 9 (Roche) and 500ng of plasmid DNA per well. After 48 hr, cells were rinsed once with PBS before being treated with Trypsin-EDTA (Gibco) for 5 min to obtain a single-cell suspension.

### Antibody staining

Cell suspensions were transferred into a 96-well plate and fixed with 2% paraformaldehyde solution for 10 min. Cells were then pelleted at 1500g for 2 min, washed with Flow Cytometry Staining Buffer (FC001, R&D Systems), and resuspended in ice-cold methanol at 4°C for 30 min. Cells were washed twice with staining buffer before incubating with fluorescently-labeled antibodies in the dark for 45 min at 4°C. Cells were washed once with staining buffer and then resuspended to a final volume of 60 µL/well in the same buffer. Cells were kept on ice until samples could be analyzed by spectral flow cytometry (Cytek Aurora CS) on the same day.

Antibodies against pAkt1 (Invitrogen, Cat # 48-9715-42, used at 1:400), pERK1/2 (Invitrogen, Cat # 46-9109-42, used at 1:200), pCREB (Cell Signaling Technology (CST), Cat #14001, used at 1:400), pS6 (CST, Cat # 4858, used at 1:50), pp38 MAPK (CST, Cat # 8632, used at 1:200) and anti-HA (Invitrogen, Cat # 26183-D680, used at 1:400) were previously validated ^19; 20; 38^ and re-assessed in our system.

### Single-cell analysis with SCADR

SCADR standalone app (https://github.com/JerryTong-GH/SCADR) was created in MATLAB Version: 9.13.0 (R2022b) with statistical scripts imported from standard R packages and MATLAB libraries. The app comes with a utility for customization of data presentations. Instructions and tutorials with examples are available on GitHub with active monitoring. Briefly, FCS data files from flow cytometer were imported into the SCADR app. Post-acquisition quality control involves removing debris using Forward Scatter (FSC) and Side Scatter (SSC), removing doublets with FSC-W and finally, post-processing data analysis only include 2-98% percentile of all phosphoproteins to reduce background noise due to outliers.

Only PTEN-expressing cells were included in the analysis by setting the lower limit of HA signal intensity based on the background signal from untransfected cells. Artifactual cellular responses were minimized by removing extremely high expressors (top 2% of HA expression) that were likely to be to induce phenotypes that are beyond the physiological level. Fluorescence-minus-one (FMO) controls ^39^ were used to reduce artificial overlap between each fluorescent marker. The number of single cells for each variant after the pre-processing was in the range of 1000-3000 per replicate depending on the transfection efficiency of each construct.

Following preprocessing, single cell signaling data were analyzed in SCADR using a suite of quantitative and multidimensional features. Initial comparisons were performed using whole population mean fluorescence intensity (MFI) values for each phosphoprotein. To capture variant expression dependent effects, cells were subsequently separated into 10 equal width expression bins based on HA PTEN levels, enabling bin wise quantification of signaling responses and statistical comparisons across variants. Dual marker dose response scatter plots were generated to visualize relationships between signal protein activity states and reveal variant specific patterns not apparent from single marker analyses. Pairwise phosphoprotein level correlations were computed to assess pathway co regulation and identify coordinated signaling modules. For higher dimensional visualization, UMAP ^40^ embeddings were constructed using all five phosphoprotein readouts, allowing exploration of variant driven cellular states across low and high expression subsets. Finally, hierarchical agglomerative clustering (AGNES) ^41^ was applied both to PTEN variants and to the phospho markers, providing an unbiased assessment of similarity relationships and revealing functional groupings within variants and signaling pathways.

### Statistical analysis

Statistical analyses were performed within the SCADR framework using standard MATLAB and Python statistical libraries. Differences in phosphoprotein signaling across PTEN variants were assessed using one-way ANOVA applied independently within each HA-expression bin. ANOVA p-values were used to determine whether variant means differed significantly, and post-hoc pairwise comparisons were visualized as heatmaps to highlight bin-specific divergence among variants. For multidimensional analyses, pairwise Pearson correlations were computed between phospho-markers to quantify co-regulation and pathway coupling. Dual-marker dose-response scatter plots were used to visualize relationships between signaling readouts, while expression-binned analyses enabled detection of expression-dependent effects that are obscured in whole-population MFI measurements. UMAP embeddings were generated from all five phosphoprotein features to capture nonlinear structure in single-cell signaling states, and the AGNES hierarchical clustering algorithm was applied to both variants and markers to identify similarity relationships and functional groupings.

## RESULTS

### SCADR analytic software for discriminating variant impacts on signaling pathways

SCADR allows input of cytometry data sets in .CSV file format from conventional and spectral flow cytometry, as well as CyTOF ^42^ (**Fig. 1A**). Dead cells and debris can be excluded from analyses with polygon gating of scatter plots of individual cell measures or with frequency histograms (**Fig. 1B**). While SCADR can perform conventional metrics, such as population-based median fluorescence intensity (MFI), SCADR is optimized for conducting analytics that preserve single-cell measures, or that bin cells based on expression levels of selected protein markers (**Fig. 1C**). SCADR is particularly effective at characterizing dose-dependent effects of exogenously-expressed protein variants, detected using conjugated antibodies against a synthetic protein tag, on the activity states of signaling proteins, via quantifying fluorescence intensities of tagged phospho-antibodies. Dose-dependent effects can be examined by plotting single-cell fluorescence measures of signal protein phosphorylation states against exogenous protein variant levels, and fitting values using regression algorithms including rLOESS (Robust Locally Estimated Scatterplot smoothing) with weighted smoothing curve fitting ^43^. The slope of fitted lines indicates the positive or negative interaction between the expression level of exogenous protein and each signaling protein, and strength of interactions. Multi-phasic dose-dependent effects can be statistically analyzed by binning cells by windows of exogenous protein variant expression, or levels of any selected endogenous marker, using user-defined bin sizes. SCADR also allows correlation analyses between levels of phosphoprotein activation states based on level of variant protein expression, or independent of variant expression. Correlation measures can indicate whether phosphoproteins interact and whether these interactions are differentially impacted by missense variants, and variant expression levels. Further, dimension reduction and clustering approaches are available to assist in identifying interactions between signaling proteins, and for clustering variants based on functional consequences. When testing large numbers of variants, clustering allows the identification of a smaller set of representatives of distinct dysfunctional classes to move forward in lower-throughput assays.

**Figure 1:**
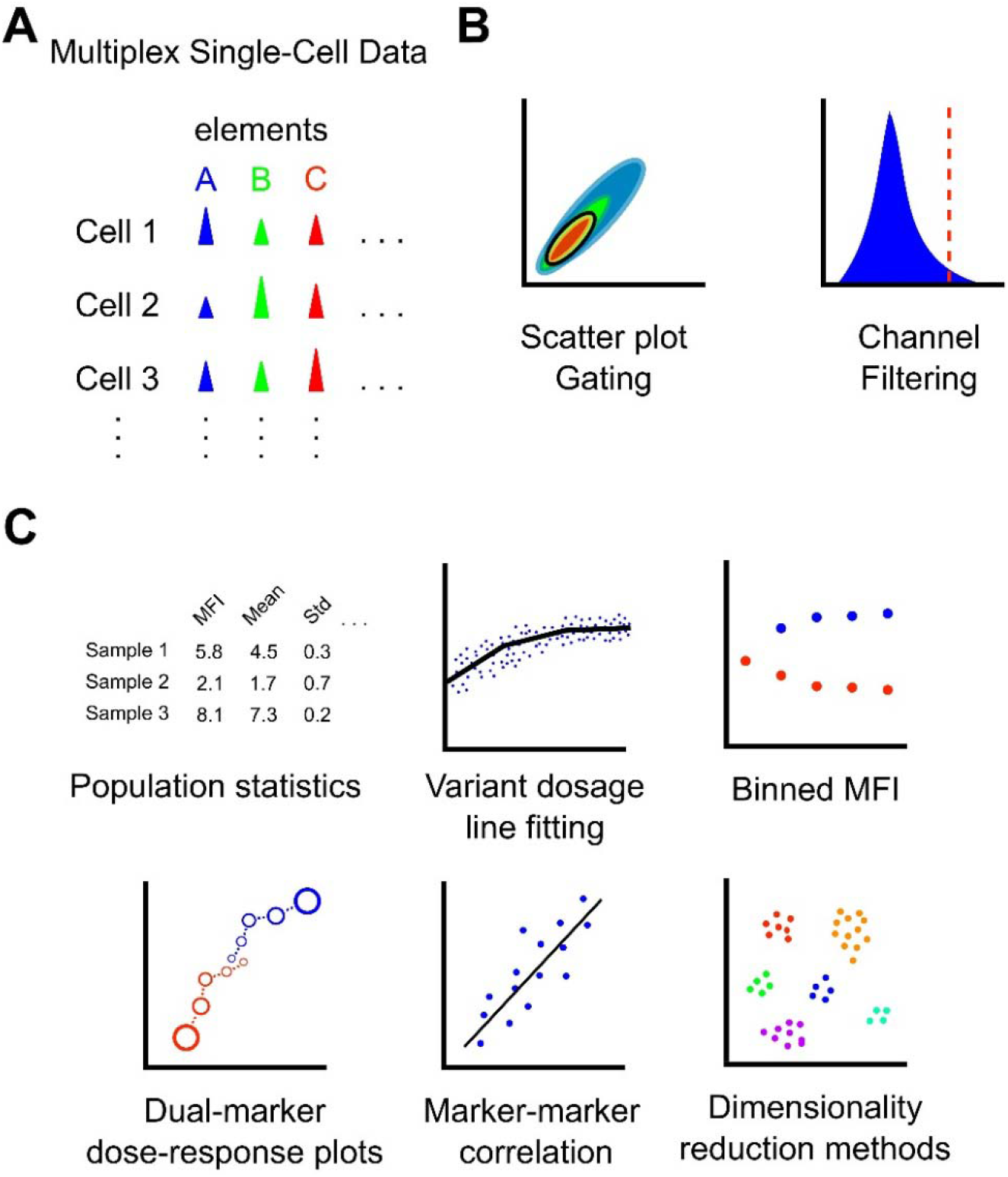
SCADR population-based, single-cell, and dose-response analytics. **(A)** SCADR is compatible with various single-cell technologies, including conventional and spectral flow cytometry, and CyTOF. **(B)** After files have been loaded into SCADR, single-cell data can be processed to remove debris and dead cells using scatter plot gating and channel filtering. **(C)** SCADR contains multiple analytical tools, including population-based measurements such as median fluorescence intensity (MFI); dosage analytics, including plotting single-cell measures against variant expression levels, and fitting distributions using linear regression and multi-exponential segmentation, binned MFI based on selected marker expression levels, dual-marker dose-response plots, and correlation analyses and dimensionality reduction methods (PCA, t-SNE, UMAP, AGNES) for clustering of variants and markers.

### Single-cell analyses of PTEN variant impacts on phosphoprotein signaling

To demonstrate the utility of SCADR for examining the impact of PTEN variants on downstream signaling cascades, we created a plasmid expression vector encoding the NCBI Reference Sequence of PTEN (RefSeq) with an N-terminal HA-tag driven by a CMV promoter. Site-directed mutagenesis was used to create 6 variants of PTEN, including the well-characterized gain-of-function (GoF) variant 4A, generated by replacing the serine and threonine phosphorylation sites (Ser380, Thr382, Thr383, and Ser385) with alanine, and the loss-of-function (LoF) variants C124S (lipid and protein phosphatase null), G129E (lipid phosphatase null), and Y138L (protein phosphatase null) (**Fig. 2A**). We also included 2 less well-characterized variants found in the human population, including P38H, first identified in an individual with ASD ^44^ and reported to be lipid phosphatase null ^20^, and D268E, found in the general population and reported to be indistinguishable from RefSeq in a pAkt assay ^20^.

**Figure 2:**
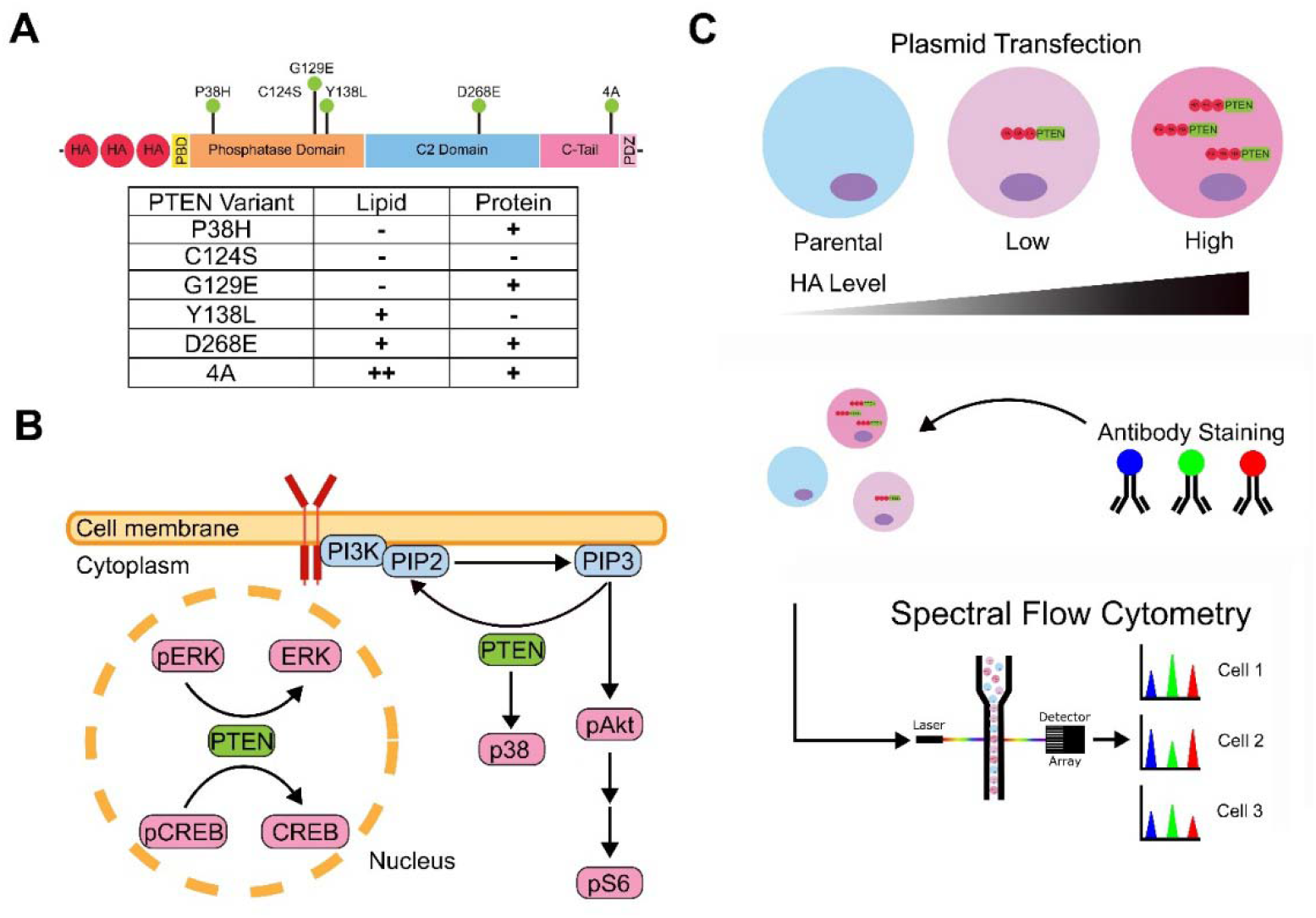
Experimental workflow for assessing impact of PTEN missense variants on canonical and noncanonical signaling using spectral flow cytometry. **(A)** A schematic diagram showing major domains of PTEN with a 3XHA at its N-terminal and sites of variants. PBD = phospholipid binding domain. Table lists PTEN variants tested in addition to the RefSeq control with their reported lipid and protein phosphatase activities: absent (-), present (+), enhanced gain-of-function (++). **(B)** A simplified PTEN signaling pathways focusing on the phosphoprotein markers used in this study. PTEN is a negative regulator of PI3K/Akt/mTOR signaling through its lipid phosphatase activity, hydrolyzing PI(3,4,5)P3 (PIP3) to PI(4,5)P2 (PIP2) leading to dephosphorylation and inhibition of Akt kinase and downstream S6 kinase activity. PTEN also functions as a protein phosphatase regulating ERK and CREB signaling in the nucleus. Reports implicate p38 phosphoprotein crosstalk with ERK and Akt pathways, as well as p38 acting upstream of PTEN. **(C)** Schematic summarizing the experimental workflow of spectral flow cytometry. HEK293 cells transfected for expression of HA-tagged PTEN variants were stained with fluorescently-conjugated antibodies and single-cell fluorescent profiles recorded.

We selected 5 phosphoproteins in signaling pathways downstream of PTEN, and corresponding phospho-antibodies sensitive to the activity states of each to measure impact of PTEN variant expression on both canonical and noncanonical pathways (**Fig. 2**). These include a phospho-antibody targeting Ser473 on Akt, as PTEN-mediated downregulation of Akt activity detected by reduced levels of phosphorylated pAkt is considered the gold standard for determining pathogenicity of PTEN variants ^33; 45^. To distinguish the impact of PTEN variants on proximal and distal signaling targets, we examined pS6, a downstream target of Akt, using an antibody targeting S6 Ser235/Ser236 ^46^. We also selected target proteins in noncanonical pathways, including PTEN’s recently described Akt-independent roles in nuclear signaling via regulating the levels of pERK1/2, detected with an antibody targeting Thr202, Tyr204 ^47; 48^, and pCREB, detected with an antibody targeting Ser133 ^36^ (**Fig. 2B**). Further, given the presence of cross-talk between PI3K/Akt/mTOR and MAPK/ERK pathways ^49^, we also included p38, a MAPK that has been proposed to act downstream of ERK, as well as Akt, using an antibody against Thr180/Tyr182 of p38 ^32; 50^.

Human embryonic kidney (HEK293) cells were transfected with each PTEN variant, and after 48 hr were fixed and labeled with fluorophore-conjugated antibodies recognizing activity states of the 5 phosphoproteins and expression levels of PTEN variants by detecting their HA-tag. Antibody staining levels were measured using a Cytek Aurora Spectral Flow Cytometer with 64 fluorophore emission detection windows (15-34 nm domains) spanning wavelengths from 365-829 nm. Spectral flow cytometry uses fluorescence detected across this large number of sampling windows to discriminate multiple fluorophores by mathematically separating the known emission fluorescence spectral signatures of each fluorophore ^51; 52^. A key feature of our multiplex analysis is the ability to measure abundance of phosphoproteins relative to the level of exogenous PTEN variant expression in each cell by tracking the intensity of a fluorophore-conjugated antibody recognizing the HA-tag of each variant (**Fig. 2C**). Since transient transfection of plasmid DNA produces cells with a wide range of plasmid load, and subsequent variant protein expression, this approach allows study of the dose-response relationships between PTEN variant protein levels and signaling protein activity states, and facilitates revealing cross-talk in signal pathway interactions.

### SCADR analyses of PTEN variant impacts on cellular signaling

#### Whole cell population MFI

Within each experiment, transfected HEK293 cells were selected using gating based on detectable levels of exogenous PTEN variant protein expression with HA signals. We first employed SCADR to conduct conventional whole cell population-based MFI measures of 5 signaling phosphoprotein activity states in cells expressing the control PTEN RefSeq and 6 PTEN variants 2 days following transfection (**Fig. 3**). Examining the impact of PTEN variants on pAkt levels revealed distinct effects of the lipid phosphatase-null mutants C124S, G129E, and P38H. All three showed significantly elevated pAkt levels compared to RefSeq, consistent with their lack of lipid phosphatase activity and dominant negative effects ^20; 32^ (**Fig. 3A**). Interestingly, the protein phosphatase null mutant Y138L also exhibited increased pAkt compared to RefSeq, although to a lesser degree, suggesting either partial deficiency in lipid phosphatase activity, or contribution of protein phosphatase activity to regulation of pAkt. In contrast, the PTEN variant D268E showed significant reductions in pAkt levels compared to RefSeq, demonstrating GoF, and was not significantly different from 4A. 4A exhibited reduced pAkt levels, but was not significantly different from RefSeq. Phosphorylation patterns of S6 following PTEN variant expression were similar to pAkt, showing significant increase in pS6 levels in cells expressing the lipid phosphatase-null mutants C124S, G129E, and P38H compared to RefSeq, D268E, 4A and Y138L (**Fig. 3B**). For pp38 and pERK, we found no detectable differences between variants (**Fig. 3C,D**). For pCREB, all variants except G129E significantly reduced pCREB levels compared to RefSeq (**Fig. 3E**), while, D268E and G129E, and P38H and G129E, were significantly different from each other.

**Figure 3:**
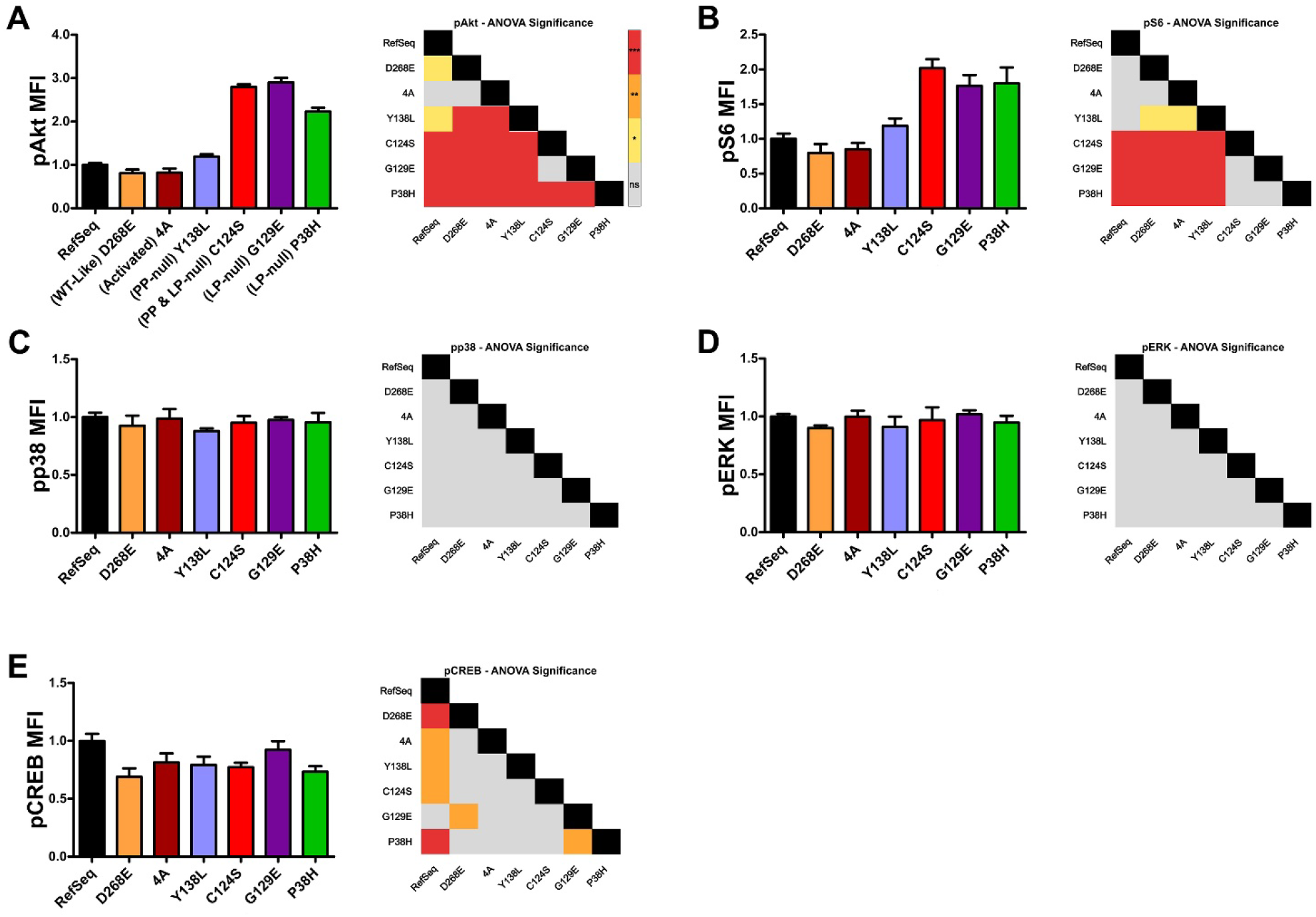
Population-based analysis of PTEN pathway phospho-markers using SCADR. **(A-E)** Whole cell population median fluorescence intensities (MFI) of pAkt, pS6, pp38, pERK and pCREB for each PTEN variant. Each MFI is the average of 4 independent wells and normalized to RefSeq under basal conditions. Statistics using a one-way ANOVA were analyzed and presented as a heatmap to the right of each histogram (Tukey post-hoc test, ns = not significant, *grey*; *p<0.05, *yellow*; **p<0.01, *orange*; ***p<0.001, *red*).

#### Single-cell relationships between PTEN variant expression level and signal protein activity

Using the same datasets, we explored the complex dose-dependent relationships between PTEN variants and associated signaling activity using SCADR (**Fig. 4A**). Linear regression analysis revealed consistent downregulation of pAkt, which decreased with increased variant PTEN expression for D268E, 4A and Y138L, to reach a similar level as with expression of RefSeq, but exhibiting different rates of decrement (**Fig. 4B**). Notably, both 4A and D268E demonstrated greater suppression of pAkt at lower expression levels than RefSeq, consistent with GoF phenotypes. In contrast, C124S, G129E and P38H all exhibited enhanced levels of pAkt, which increased as PTEN variant expression increased ^44^ (**Fig. 4B**).

**Figure 4:**
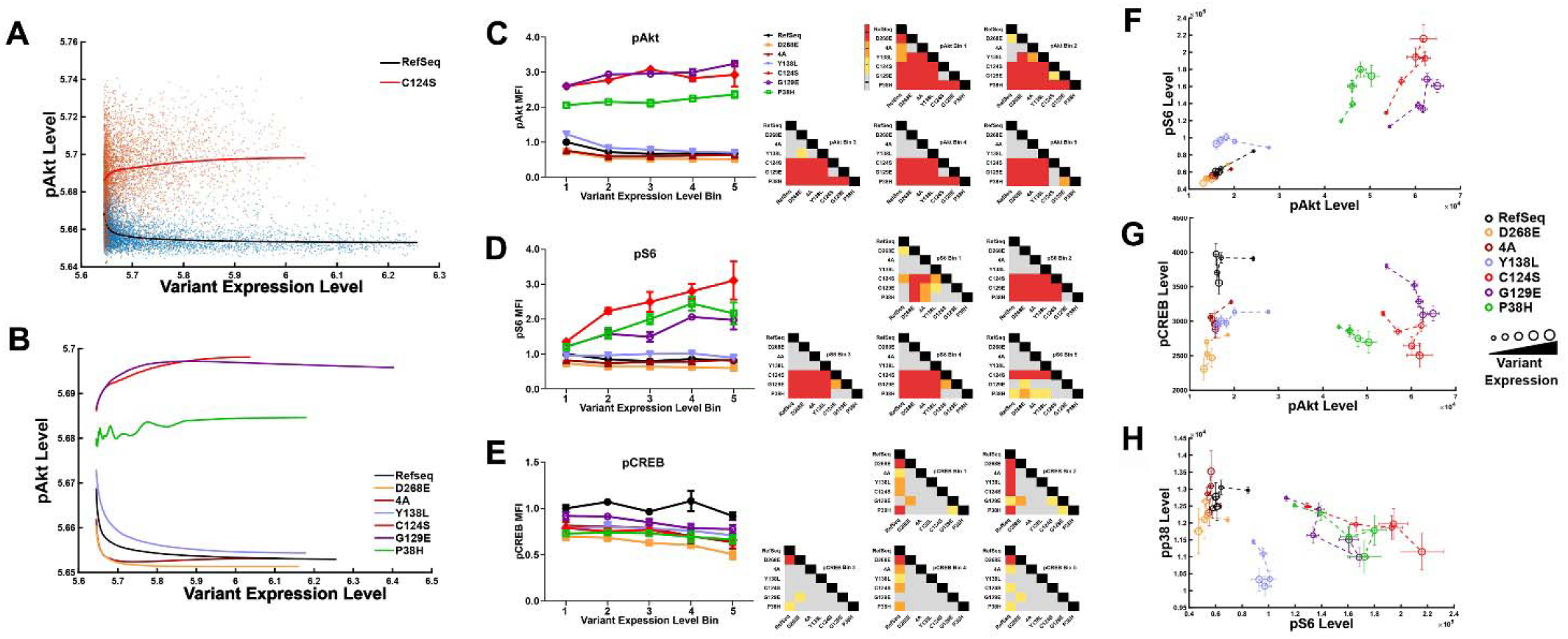
Dose-response analyses by SCADR reveals subtle differences in regulation of phosphoproteins by PTEN variants. **(A)** Scatter plots of single-cell fluorescence measures showing the expression levels of pAkt in PTEN RefSeq and PTEN C124S expressing cells plotted against HA intensities indicating expression level of exogenous PTEN variants in each cell. Solid lines show the rLOESS (Locally Estimated Scatterplot Smoothing) weighted smoothing regression of single-cell data points. **(B)** pAkt LOESS regression curves for all variants. **(C-E)** All cells are grouped into 10 bins based on exogenously expressed protein abundance (HA intensities), and the MFI values of the first 5 bins are plotted for phosphoproteins pAkt, pS6 and pCREB with significance differences between all variants measured using one-way ANOVA and presented as a heatmap to the right of each histogram (Tukey post-hoc test, ns = not significant, *grey*; *p<0.05, *yellow*; **p<0.01, *orange*; ***p<0.001, *red*). **(F-H)** Dual-marker dose-response curves using the first 5 expression level bins, with increasing variant expression level indicated by increasing circle size, demonstrating variant-specific interactions between pairs of phosphoproteins with increasing expression levels of each PTEN variant.

To further quantify these effects, we grouped single-cell data into 10 bins of equal width based on PTEN variant expression levels (**Fig. 4C-E**). Since missense variants can impact protein abundance ^19; 20; 53^, we focused on moderate expression bins (bins 1-5), which contained consistently high cell numbers across all tested variants for comparing their impacts on signaling proteins. Single-cell analytics indicate that whole cell population MFI values are heavily weighted by cells with low variant expression and that binning based on expression reveals more complex impacts on signaling proteins. At low variant expression, pAkt modulation was significantly different from RefSeq for all variants, and all variants exhibited significantly different values except the pairs 4A and D268E, and C124S and G129E. As variant expression levels increase, D268E, 4A and Y138L become indistinguishable from RefSeq and each other, and C124S becomes indistinguishable from P38H. pS6 activity patterns show similar results with variants retaining lipid phosphatase activity (D268E, 4A, Y138L) being significantly different from lipid phosphatase-null variants (C124S, G129E, P38H) across most bins. Interestingly, dose-response measures identified significant differences between the 3 lipid phosphatase-null mutants not seen with whole cell population MFI. The binned dose response data clearly demonstrated differences between P38H, C124S, and G129E for pAkt and pS6, indicating overlapping, but distinct dysfunctional impacts. For pCREB, binned dose response values (**Fig. 4E**) largely matched whole cell population MFI, with the exception of significant difference between C124S and G129E observed at moderate variant expression levels. Overall, results demonstrated the utility of SCADR to discriminate subtle but consequential impacts on signaling cascades by PTEN variants that are lost in aggregated MFI analyses.

To visualize how activity states of pairs of signaling proteins correlate as PTEN variant concentrations change, SCADR displays dual-marker dose-response plots with cells binned by variant expression level with increasing levels represented by larger circle sizes (**Fig. 4F-H**). These two-dimensional projections provide a visualization of how signal proteins interact at different cell states, which can reveal distinctions between variants not apparent from single-marker measures. In the dual-marker dose-response plots pS6 vs. pAkt, pCREB vs. pAkt, and pp38 vs. pS6 (**Fig. 4F-H**), variants with and without lipid phosphatase activity are well segregated, and variants within these groups have unique profiles. For example, the Y138L profile is similar to, but well separated from D268E, 4A, and RefSeq, and P38H is distinct from C124S and G129E. Together, these dual marker dose-response plots demonstrate how subtle differences in PTEN’s lipid and protein phosphatase functions generate distinct multidimensional signaling landscapes, and how specific mutations reshape pathway cross talk to produce variant specific signaling phenotypes.

#### SCADR correlation analyses reveal PTEN variant-dependent cross-talk between phosphoproteins

To further investigate interactions between signaling proteins and how these relationships are impacted by PTEN variants, independent or dependent on variant expression levels, we used SCADR to compute Spearman correlation coefficients (r) between signal protein activities across all single-cell measures, as well as within cells with low or high PTEN variant expression (**Fig. 5A-C**). Correlation analyses discriminated variants with lipid phosphatase function and dysfunction, and showed both expected interactions within canonical pathways, as well as cross-talk with and within noncanonical pathways. As in the dual-marker dose-response plots, we found strong positive correlations between pAkt and pS6 in cells expressing variants with lipid phosphatase activity (RefSeq, D268E, 4A, Y138L) due to strong inhibition of this pathway. Y138L showed the lowest correlation within this cohort, supporting a role for protein phosphatase activity in pAkt/S6 signaling. Weaker correlations between pAkt and pS6 were observed in lipid phosphatase-deficient variants (C124S, G129E, P38H). We found little impact of PTEN variants on ERK activity and corresponding low correlations between ERK with other signaling proteins in either canonical or noncanonical pathways. Interestingly, we observed a strong correlation between pp38 and pAkt/pS6 in cells expressing variants with intact lipid phosphatase activity, but not lipid phosphatase-deficient variants, which is heightened in cells with higher exogenous variant expression, supporting previous reports of a p38/PTEN/Akt signaling axis ^54^. When comparing across variants, we observed grouping patterns based on their catalytic functionality. RefSeq, D268E, 4A and to lesser degree Y138L showed similar impacts on paired phosphoprotein activity correlations, with these variants associated with a positive correlation in pAkt/pS6, pAkt/pCREB, pAkt/pp38, and pS6/pp38. These results underscore the likelihood of PTEN’s lipid and protein phosphatase activities in co-mediating cellular signaling and the utility of dose-dependent analysis for classification of variant functions.

**Figure 5:**
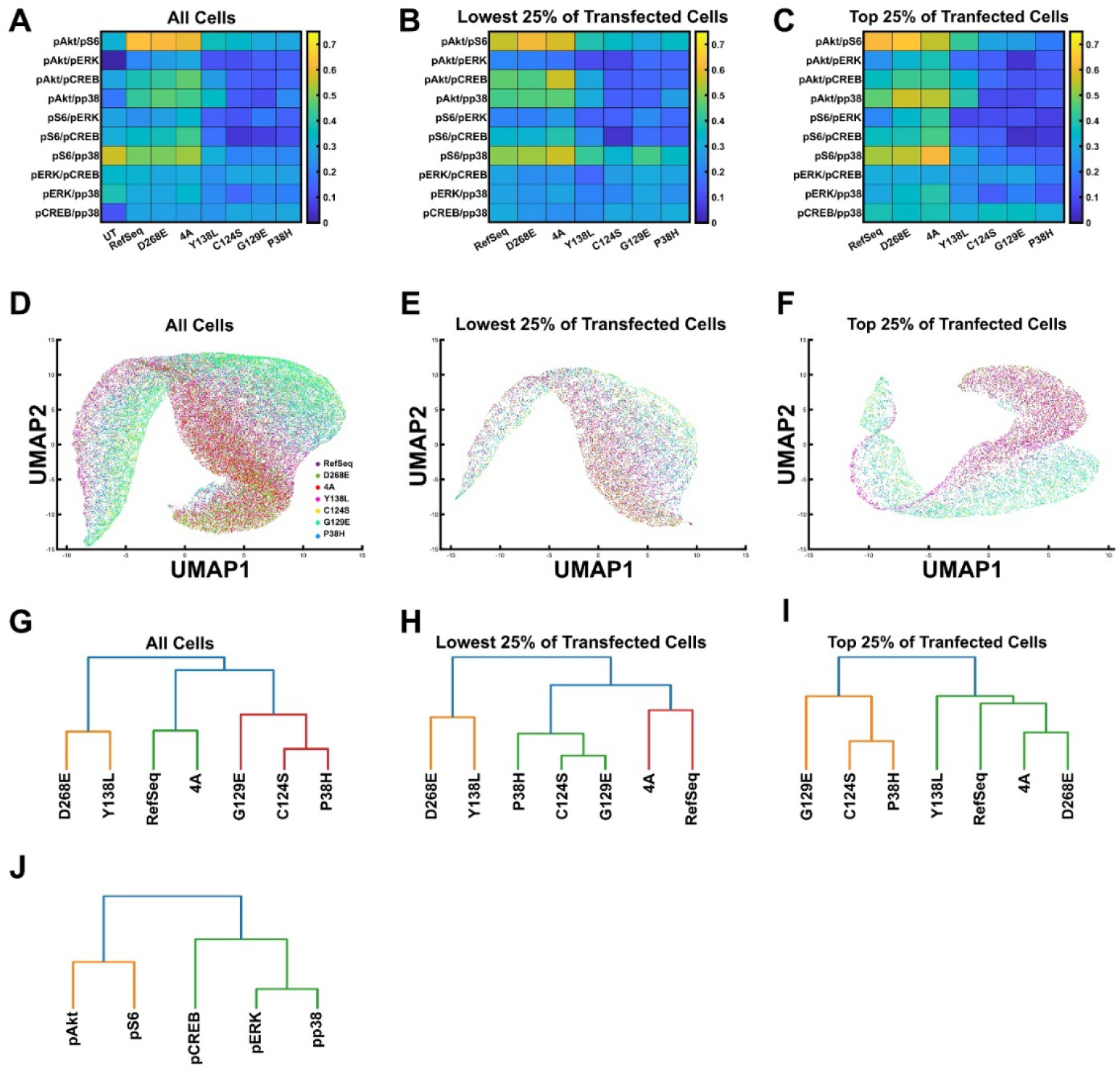
SCADR single-cell analyses reveal PTEN variant-dependent signaling cross-talk and functional clustering. Spearman correlations of phosphoprotein activity states across PTEN variants reveal distinct impacts on signaling relationships. **(A)** Pairwise correlation heatmap of phosphoprotein activity states across all cells expressing PTEN variants compared to untransfected cells (UT). **(B)** Pairwise correlation of cells with the lowest quartile of exogenous PTEN variant expression. **(C)** Pairwise correlation of cells with the top quartile of PTEN variant expression. (**D-F**) UMAP dimensionality reduction of single-cell phosphoprotein profiles distinguishes variant-driven cellular states in all cells (**D**), cells with the lowest (**E**) and highest (**F**) quartile of exogenous PTEN variant expression. The dimensions pAkt, pS6, pp38, pCREB and pERK were reduced to 2 dimensions (n-neighbors: 75, min-distance: 0.6). **(G-I)** AGNES hierarchical clustering of PTEN variants based on five phospho-marker profiles in all cells (**G**), cells with the lowest (**H**) and highest (**I**) quartile of exogenous PTEN variant expression. **(J)** AGNES hierarchical clustering of the five phospho markers (pAkt, pS6, pERK, pCREB, pp38) in all transfected cells.

### Dimensional reduction and clustering distinguish signal protein relationships and variant impacts

SCADR includes multiple dimensionality reduction methods, including Principal Component Analysis (PCA) ^55^, t-distributed stochastic neighbor embedding (t-SNE) ^56^, and Uniform Manifold Approximation Projection (UMAP). Using UMAP ^40^, we visualized single-cell distribution across PTEN variants in a HA-independent manner (**Fig. 5D-F**). Individual cell expression levels of the five phosphoproteins pAkt, pS6, pp38, pERK, and pCREB were used to generate the UMAP embedding. This spatial distribution of signaling profiles suggests distinct variant-driven cellular states, likely influenced by variant expression levels. The overall pattern was consistent when restricting analysis to cells expressing low levels of exogenous PTEN variants, but with less separation (**Fig. 5E**). Cells with higher expression levels exhibited sharper separation of variants into two well-defined groups: RefSeq, D268E, 4A and Y138L; and another comprising C124S, G129E and P38H, indicating context-dependent functionality of individual variants (**Fig. 5F**).

To evaluate how individual PTEN variants relate to one another across the five functional markers measured, we performed AGNES algorithm-based hierarchical agglomerative clustering^57^. This analysis revealed clear similarity relationships among variants. As expected, the lipid phosphatase–deficient variants (G129E, C124S, and P38H) clustered together (**Fig. 5G**). Notably, P38H grouped closely with C124S, supporting its classification as a lipid phosphatase–null variant. In contrast, D268E and Y138L formed a distinct cluster, indicating that D268E may also exhibit reduced protein phosphatase function despite its previously described RefSeq-like behavior. To assess how PTEN variant dosage influences hierarchical clustering of downstream signaling, we compared clustering patterns across all transfected cells, the lowest 25% of PTEN expressing cells, and the highest 25% of PTEN expressing cells using phospho signaling markers (**Fig. 5G–I**). When analyzing the lowest 25% of transfected cells, the clustering pattern closely mirrored the structure observed when all cells were included (**Fig. 5H**). Lipid phosphatase–null variants continued to group together. In contrast, the highest 25% of transfected cells revealed a distinct clustering state (**Fig. 5I**). While lipid phosphatase–null mutants still grouped together as expected, a new cluster emerged containing Y138L, RefSeq, 4A, and D268E. Within this cluster, Y138L formed the most divergent branch, whereas the activated variants 4A and D268E clustered together. These shifts in cluster structure at high PTEN expression levels highlight a dosage dependent reorganization of PTEN regulated signaling. Together, these analyses demonstrate that PTEN variant dosage can reshape the signaling landscape, revealing variant specific behaviors that are not apparent when all cells are analyzed collectively. We also applied AGNES hierarchical clustering ^57^ to the 5 phospho-markers (**Fig. 5J**), showing the expected pAkt/pS6 tightest cluster, consistent with their shared dependence on PI3K/Akt/mTOR signaling and their role as primary readouts of PTEN lipid-phosphatase activity. A second cluster comprised pERK and pp38, reflecting coordinated behavior within the MAPK pathway. pCREB occupied an intermediate position, showing greater similarity to the MAPK-associated markers but remaining distinct from both major clusters. This pattern suggests that pCREB integrates inputs from multiple pathways and may capture signaling effects not fully represented by either the Akt/S6 or ERK/p38 modules alone.

## DISCUSSION

Many high-throughput approaches for quantifying protein expression or phosphorylation state, such as those based on a western blotting platforms and protein lysate microarrays combine samples from large populations of cells ^58–60^. Although single-cell technologies such as conventional or spectral flow cytometry and CyTOF record single-cell measures, it is standard practice to aggregate data into population metrics. However, while aggregation is used, in part, to compensate for cellular heterogeneity, preserving single-cell measures in large population datasets allows correlating phenotypes with distinct cell states. Single-cell analytics can take advantage of ranking cells based on expression levels of endogenous or exogenous protein markers for investigating dose-response relationships. This is particularly advantageous in experiments using exogenous protein expression through plasmid-based transfection, which inherently yields a wide range of expression levels due to variable plasmid delivery to each cell. While such variability can be a disadvantage for population aggregation approaches, it offers a rich dataset for investigating complex dose-response effects that may discriminate subtle impacts such as those induced by different missense variants of the same protein. Multiplex measures of the activity states of distinct and overlapping signaling pathways in the same cells allow interrogation of the types and levels of dysfunction induced by missense variants. Results can determine whether variant-induced dysfunctions affect all or a subset of protein functions, and can elucidate potential roles of canonical and noncanonical pathways in pathophysiology. Although flow cytometry measures of fixed cells are static, multiplex measures from large numbers of cells allow characterization of dynamic signal protein interactions based on the range of physiological states in the cell population sampled. Our approach contrasts with other single-cell cytometry algorithms ^61; 62^ that are optimized mainly to identify distinct cell populations based on the presence of biomarkers or changes in their levels, but is distinguished by its focus on dose-response relationships.

Here, we compare population and single-cell analytics using our custom-designed software SCADR applied to a multiplex spectral flow cytometry dataset to distinguish impacts of 6 PTEN missense variants on activity states of 5 phosphoproteins in canonical and noncanonical downstream signaling pathways. Results indicate variant-specific effects on both canonical and noncanonical PTEN signaling pathways. Comparison of interactions between signal proteins showed high correlation between pAkt and pS6, confirming that both proteins are in the same Akt/mTOR pathway ^33^; ^34; 46^. Dephosphorylation of Akt observed at lower PTEN expression levels than S6 supports S6 being downstream of Akt. Correlations between the activity states of Akt, CREB, and p38, and between S6 and p38 were enhanced in cells expressing PTEN variants with lipid phosphatase function, indicating a role for PTEN in regulating both CREB and pp38 within the Akt/S6 pathway ^36; 47; 48^.

SCADR is particularly useful for characterizing and discriminating the impact of different missense variants on multiple protein functions. Critically, no two variants had identical dose-sensitive impacts on the signal proteins examined. In particular, results distinguish the impacts of the variants Y138L, G129E, and D268E. Results with Y138L support a role for PTEN protein phosphatase activity in regulation of pp38. G129E exhibited similar impact on pAkt as C124S, but differed in its effects on pS6. Further, SCADR distinguishes differences between the lipid phosphatase LoF variants G129E and P38H on pAkt and pS6. Moreover, SCADR results found that D268E, a variant previously reported to be indistinguishable from RefSeq, exhibited significant differences in impact on both canonical and noncanonical signaling.

Overall, results demonstrate the complexity of different missense variant impacts on multiple protein functions, countering common models that attribute pathogenic missense variant impacts to haploinsufficiency mediated by a general reduction in all protein functions due to protein instability and loss. Rather, restricted functional impacts can occur independent of effects on protein stability or expression levels ^20^. Since single-cell analytics with SCADR allows tracking multiple impacts based on levels of variant expression, one can distinguish between haploinsufficiency and more complex effects on distinct protein functions. SCADR coupled to multiparameter cytometry expedites the process of mapping large numbers of gene variants with unknown clinical impact to molecular dysfunctions and the phenotypic spectrum of disease outcomes.

## ACKNOWLEDGMENTS

This work has been funded by the PTEN Research Foundation, a charity governed by English law (charity number 117358) to Dr Kurt Haas under grant number UBC-21-001.

## AUTHOR CONTRIBUTIONS

C.G. and M.T. contributed equally to this work.

Conceptualization, K.H.; Formal Analysis, C.G., W.C.S., W.M.M., and J.S.T.; Funding Acquisition, K.H.; Investigation, C.G., W.C.S., F.M., and W.M.M. [ADD M.T. HERE IF APPLICABLE]; Methodology, J.S.T., Y.H., C.G., W.C.S., F.M., and W.M.M.; Project Administration, K.H.; Software, J.S.T., Y.H., and P.C.; Supervision, K.H. and P.P.; Writing – Original Draft, C.G. and K.H.; Writing – Review & Editing, C.G., W.C.S., W.M.M., J.S.T., and K.H.

## WEB RESOURCES

The data presented in this manuscript were generated by the authors and available on request. We have developed a scalable open-source software platform for multiplex sampling of canonical and noncanonical signaling pathways called **S**ingle-**C**ell **A**nalytics for **D**ose-**R**esponse (**SCADR**), available at: github.com/JerryTong-GH/SCADR. SCADR is freely available for download through GitHub. The software includes a fully developed user interface, enabling researchers with varying levels of computational experience to use the platform. The GitHub repository also provides comprehensive documentation describing all software functionalities, along with automated MATLAB-based testing procedures to ensure code integrity and maintain software reliability during future updates.

## REFERENCES

1. Genomes Project, C., Auton, A., Brooks, L.D., Durbin, R.M., Garrison, E.P., Kang, H.M., Korbel, J.O., Marchini, J.L., McCarthy, S., McVean, G.A., et al. (2015). A global reference for human genetic variation. Nature 526, 68–74.

2. Lek, M., Karczewski, K.J., Minikel, E.V., Samocha, K.E., Banks, E., Fennell, T., O’Donnell-Luria, A.H., Ware, J.S., Hill, A.J., Cummings, B.B., et al. (2016). Analysis of protein-coding genetic variation in 60,706 humans. Nature 536, 285–291.

3. All of Us Research Program, I., Denny, J.C., Rutter, J.L., Goldstein, D.B., Philippakis, A., Smoller, J.W., Jenkins, G., and Dishman, E. (2019). The “All of Us” Research Program. N Engl J Med 381, 668–676.

4. Sudlow, C., Gallacher, J., Allen, N., Beral, V., Burton, P., Danesh, J., Downey, P., Elliott, P., Green, J., Landray, M., et al. (2015). UK biobank: an open access resource for identifying the causes of a wide range of complex diseases of middle and old age. PLoS Med 12, e1001779.

5. Do, R., Kathiresan, S., and Abecasis, G.R. (2012). Exome sequencing and complex disease: practical aspects of rare variant association studies. Hum Mol Genet 21, R1–9.

6. Laddach, A., Ng, J.C.F., and Fraternali, F. (2021). Pathogenic missense protein variants affect different functional pathways and proteomic features than healthy population variants. PLoS Biol 19, e3001207.

7. Sahni, N., Yi, S., Taipale, M., Fuxman Bass, J.I., Coulombe-Huntington, J., Yang, F., Peng, J., Weile, J., Karras, G.I., Wang, Y., et al. (2015). Widespread macromolecular interaction perturbations in human genetic disorders. Cell 161, 647–660.

8. Stenson, P.D., Mort, M., Ball, E.V., Shaw, K., Phillips, A., and Cooper, D.N. (2014). The Human Gene Mutation Database: building a comprehensive mutation repository for clinical and molecular genetics, diagnostic testing and personalized genomic medicine. Hum Genet 133, 1–9.

9. Stone, E.A., and Sidow, A. (2005). Physicochemical constraint violation by missense substitutions mediates impairment of protein function and disease severity. Genome Res 15, 978–986.

10. Iossifov, I., O’Roak, B.J., Sanders, S.J., Ronemus, M., Krumm, N., Levy, D., Stessman, H.A., Witherspoon, K.T., Vives, L., Patterson, K.E., et al. (2014). The contribution of de novo coding mutations to autism spectrum disorder. Nature 515, 216–221.

11. Satterstrom, F.K., Kosmicki, J.A., Wang, J., Breen, M.S., De Rubeis, S., An, J.Y., Peng, M., Collins, R., Grove, J., Klei, L., et al. (2020). Large-Scale Exome Sequencing Study Implicates Both Developmental and Functional Changes in the Neurobiology of Autism. Cell 180, 568–584 e523.

12. O’Roak, B.J., Stessman, H.A., Boyle, E.A., Witherspoon, K.T., Martin, B., Lee, C., Vives, L., Baker, C., Hiatt, J.B., Nickerson, D.A., et al. (2014). Recurrent de novo mutations implicate novel genes underlying simplex autism risk. Nat Commun 5, 5595.

13. Vogelstein, B., Papadopoulos, N., Velculescu, V.E., Zhou, S., Diaz, L.A., Jr., and Kinzler, K.W. (2013). Cancer genome landscapes. Science 339, 1546–1558.

14. Lawrence, M.S., Stojanov, P., Polak, P., Kryukov, G.V., Cibulskis, K., Sivachenko, A., Carter, S.L., Stewart, C., Mermel, C.H., Roberts, S.A., et al. (2013). Mutational heterogeneity in cancer and the search for new cancer-associated genes. Nature 499, 214–218.

15. Cheng, J., Novati, G., Pan, J., Bycroft, C., Zemgulyte, A., Applebaum, T., Pritzel, A., Wong, L.H., Zielinski, M., Sargeant, T., et al. (2023). Accurate proteome-wide missense variant effect prediction with AlphaMissense. Science 381, eadg7492.

16. Starita, L.M., Ahituv, N., Dunham, M.J., Kitzman, J.O., Roth, F.P., Seelig, G., Shendure, J., and Fowler, D.M. (2017). Variant Interpretation: Functional Assays to the Rescue. Am J Hum Genet 101, 315–325.

17. Findlay, G.M., Daza, R.M., Martin, B., Zhang, M.D., Leith, A.P., Gasperini, M., Janizek, J.D., Huang, X., Starita, L.M., and Shendure, J. (2018). Accurate classification of BRCA1 variants with saturation genome editing. Nature 562, 217–222.

18. Fowler, D.M., and Fields, S. (2014). Deep mutational scanning: a new style of protein science. Nat Methods 11, 801–807.

19. Meili, F., Wei, W.J., Sin, W.C., Meyers, W.M., Dascalu, I., Callaghan, D.B., Rogic, S., Pavlidis, P., and Haas, K. (2021). Multi-parametric analysis of 57 SYNGAP1 variants reveal impacts on GTPase signaling, localization, and protein stability. Am J Hum Genet 108, 148–162.

20. Post, K.L., Belmadani, M., Ganguly, P., Meili, F., Dingwall, R., McDiarmid, T.A., Meyers, W.M., Herrington, C., Young, B.P., Callaghan, D.B., et al. (2020). Multi-model functionalization of disease-associated PTEN missense mutations identifies multiple molecular mechanisms underlying protein dysfunction. Nat Commun 11, 2073.

21. Fragoza, R., Das, J., Wierbowski, S.D., Liang, J., Tran, T.N., Liang, S., Beltran, J.F., Rivera-Erick, C.A., Ye, K., Wang, T.Y., et al. (2019). Extensive disruption of protein interactions by genetic variants across the allele frequency spectrum in human populations. Nat Commun 10, 4141.

22. Zaretsky, J.Z., and Wreschner, D.H. (2008). Protein multifunctionality: principles and mechanisms. Transl Oncogenomics 3, 99–136.

23. Day, E.K., Sosale, N.G., and Lazzara, M.J. (2016). Cell signaling regulation by protein phosphorylation: a multivariate, heterogeneous, and context-dependent process. Curr Opin Biotechnol 40, 185–192.

24. de Graaf, J.F., and Arens, R. (2023). Immunophenotyping beyond the limits of time. Cell Rep Methods 3, 100612.

25. George, A.A., Paz, H., Fei, F., Kirzner, J., Kim, Y.M., Heisterkamp, N., and Abdel-Azim, H. (2015). Phosphoflow-Based Evaluation of Mek Inhibitors as Small-Molecule Therapeutics for B-Cell Precursor Acute Lymphoblastic Leukemia. PLoS One 10, e0137917.

26. Toney, N.J., Schlom, J., and Donahue, R.N. (2023). Phosphoflow cytometry to assess cytokine signaling pathways in peripheral immune cells: potential for inferring immune cell function and treatment response in patients with solid tumors. J Exp Clin Cancer Res 42, 247.

27. Li, J., Yen, C., Liaw, D., Podsypanina, K., Bose, S., Wang, S.I., Puc, J., Miliaresis, C., Rodgers, L., McCombie, R., et al. (1997). PTEN, a putative protein tyrosine phosphatase gene mutated in human brain, breast, and prostate cancer. Science 275, 1943–1947.

28. Myers, M.P., Pass, I., Batty, I.H., Van der Kaay, J., Stolarov, J.P., Hemmings, B.A., Wigler, M.H., Downes, C.P., and Tonks, N.K. (1998). The lipid phosphatase activity of PTEN is critical for its tumor supressor function. Proc Natl Acad Sci U S A 95, 13513–13518.

29. Hollander, M.C., Blumenthal, G.M., and Dennis, P.A. (2011). PTEN loss in the continuum of common cancers, rare syndromes and mouse models. Nat Rev Cancer 11, 289–301.

30. Hobert, J.A., and Eng, C. (2009). PTEN hamartoma tumor syndrome: an overview. Genet Med 11, 687–694.

31. Rademacher, S., and Eickholt, B.J. (2019). PTEN in Autism and Neurodevelopmental Disorders. Cold Spring Harb Perspect Med 9.

32. Mighell, T.L., Evans-Dutson, S., and O’Roak, B.J. (2018). A Saturation Mutagenesis Approach to Understanding PTEN Lipid Phosphatase Activity and Genotype-Phenotype Relationships. Am J Hum Genet 102, 943–955.

33. Carracedo, A., and Pandolfi, P.P. (2008). The PTEN-PI3K pathway: of feedbacks and cross-talks. Oncogene 27, 5527–5541.

34. Lee, Y.R., Chen, M., and Pandolfi, P.P. (2018). The functions and regulation of the PTEN tumour suppressor: new modes and prospects. Nat Rev Mol Cell Biol 19, 547–562.

35. Song, M.S., Salmena, L., and Pandolfi, P.P. (2012). The functions and regulation of the PTEN tumour suppressor. Nat Rev Mol Cell Biol 13, 283–296.

36. Gu, T., Zhang, Z., Wang, J., Guo, J., Shen, W.H., and Yin, Y. (2011). CREB is a novel nuclear target of PTEN phosphatase. Cancer Res 71, 2821–2825.

37. Graham, F.L., Smiley, J., Russell, W.C., and Nairn, R. (1977). Characteristics of a human cell line transformed by DNA from human adenovirus type 5. J Gen Virol 36, 59–74.

38. Scurll, J.M. (2021). New methods for cluster analysis and their applications to the biology of B cells and diffuse large B-cell lymphoma. Doctor of Philosophy - PhD, University of British Columbia, Vancouver.

39. Roederer, M. (2001). Spectral compensation for flow cytometry: visualization artifacts, limitations, and caveats. Cytometry 45, 194–205.

40. McInnes, L., Healy, J., Saul, N., and Großberger, L. (2018). UMAP: Uniform Manifold Approximation and Projection. Journal of Open Source Software 3, 861.

41. Kaufman, L., and Rousseeuw, P.J. (1990). Finding Groups in Data: An Introduction to Cluster Analysis.

42. Bandura, D.R., Baranov, V.I., Ornatsky, O.I., Antonov, A., Kinach, R., Lou, X., Pavlov, S., Vorobiev, S., Dick, J.E., and Tanner, S.D. (2009). Mass cytometry: technique for real time single cell multitarget immunoassay based on inductively coupled plasma time-of-flight mass spectrometry. Anal Chem 81, 6813–6822.

43. Cleveland, W.S. (1979). Robust Locally Weighted Regression and Smoothing Scatterplots.. Journal of the American Statistical Association 74, 829–836.

44. Klein, S., Sharifi-Hannauer, P., and Martinez-Agosto, J.A. (2013). Macrocephaly as a clinical indicator of genetic subtypes in autism. Autism Res 6, 51–56.

45. Tu, T., Chen, J., Chen, L., and Stiles, B.L. (2020). Dual-Specific Protein and Lipid Phosphatase PTEN and Its Biological Functions. Cold Spring Harb Perspect Med 10.

46. Manning, B.D. (2004). Balancing Akt with S6K: implications for both metabolic diseases and tumorigenesis. J Cell Biol 167, 399–403.

47. Chung, J.H., Ostrowski, M.C., Romigh, T., Minaguchi, T., Waite, K.A., and Eng, C. (2006). The ERK1/2 pathway modulates nuclear PTEN-mediated cell cycle arrest by cyclin D1 transcriptional regulation. Hum Mol Genet 15, 2553–2559.

48. Weng, L.P., Smith, W.M., Brown, J.L., and Eng, C. (2001). PTEN inhibits insulin-stimulated MEK/MAPK activation and cell growth by blocking IRS-1 phosphorylation and IRS-1/Grb-2/Sos complex formation in a breast cancer model. Hum Mol Genet 10, 605–616.

49. Frazier, T.W., Jaini, R., Busch, R.M., Wolf, M., Sadler, T., Klaas, P., Hardan, A.Y., Martinez-Agosto, J.A., Sahin, M., Eng, C., et al. (2021). Cross-level analysis of molecular and neurobehavioral function in a prospective series of patients with germline heterozygous PTEN mutations with and without autism. Mol Autism 12, 5.

50. Xu, Y., Li, N., Xiang, R., and Sun, P. (2014). Emerging roles of the p38 MAPK and PI3K/AKT/mTOR pathways in oncogene-induced senescence. Trends Biochem Sci 39, 268–276.

51. Lazarski, C.A., and Hanley, P.J. (2024). Review of flow cytometry as a tool for cell and gene therapy. Cytotherapy 26, 103–112.

52. Sharma, S., Boyer, J., and Teyton, L. (2024). A practitioner’s view of spectral flow cytometry. Nat Methods 21, 740–743.

53. Matreyek, K.S., J.J; Ahler, E.; Fowler, D.M. (2021). Integrating thousands of PTEN variant activity and abundance measurements reveals variant subgroups and new dominant negatives in cancers. Genome Medicine 13.

54. Ming, M., Feng, L., Shea, C.R., Soltani, K., Zhao, B., Han, W., Smart, R.C., Trempus, C.S., and He, Y.Y. (2011). PTEN positively regulates UVB-induced DNA damage repair. Cancer Res 71, 5287–5295.

55. Pearson, K. (1901). LIII. On lines and planes of closest fit to systems of points in space. Philosophical Magazine Series 1 2, 559–572.

56. Downey, D., and Etzioni, O. (2008). Look Ma, no hands: analyzing the monotonic feature abstraction for text classification. In Proceedings of the 22nd International Conference on Neural Information Processing Systems. (Vancouver, British Columbia, Canada, Curran Associates Inc.), pp 393–400.

57. Pedregosa, F., Varoquaux, G., Gramfort, A., Michel, V., Thirion, B., Grisel, O., Blondel, M., Prettenhofer, P., Weiss, R., Dubourg, V., et al. (2011). Scikit-learn: Machine Learning in Python. J Mach Learn Res 12, 2825–2830.

58. Jia, Z., Barbier, L., Stuart, H., Amraei, M., Pelech, S., Dennis, J.W., Metalnikov, P., O’Donnell, P., and Nabi, I.R. (2005). Tumor cell pseudopodial protrusions. Localized signaling domains coordinating cytoskeleton remodeling, cell adhesion, glycolysis, RNA translocation, and protein translation. J Biol Chem 280, 30564–30573.

59. Komurov, K., Padron, D., Cheng, T., Roth, M., Rosenblatt, K.P., and White, M.A. (2010). Comprehensive mapping of the human kinome to epidermal growth factor receptor signaling. J Biol Chem 285, 21134–21142.

60. Treindl, F., Ruprecht, B., Beiter, Y., Schultz, S., Dottinger, A., Staebler, A., Joos, T.O., Kling, S., Poetz, O., Fehm, T., et al. (2016). A bead-based western for high-throughput cellular signal transduction analyses. Nat Commun 7, 12852.

61. Orlova, D.Y., Zimmerman, N., Meehan, S., Meehan, C., Waters, J., Ghosn, E.E., Filatenkov, A., Kolyagin, G.A., Gernez, Y., Tsuda, S., et al. (2016). Earth Mover’s Distance (EMD): A True Metric for Comparing Biomarker Expression Levels in Cell Populations. PLoS One 11, e0151859.

62. Weber, L.M., Nowicka, M., Soneson, C., and Robinson, M.D. (2019). diffcyt: Differential discovery in high-dimensional cytometry via high-resolution clustering. Commun Biol 2, 183.

